# Conserved cell-lineage controlled epigenetic regulation in human and mouse glioblastoma stem cells determines functionally distinct subgroups

**DOI:** 10.1101/2021.02.22.431953

**Authors:** Xi Lu, Naga Prathyusha Maturi, Malin Jarvius, Linxuan Zhao, Yuan Xie, E-Jean Tan, Pengwei Xing, Mårten Fryknäs, Lene Uhrbom, Xingqi Chen

## Abstract

There is ample support for developmental regulation of glioblastoma stem cells (GSCs). To examine how cell lineage controls GSC function we have performed a cross-species epigenome analysis of mouse and human GSC cultures. We have analyzed and compared the chromatin-accessibility landscape of nine mouse GSC cultures of defined cell of origin and 60 patient-derived GSC cultures by assay for transposase-accessible chromatin using sequencing (ATAC-seq). This uncovered a variability of both mouse and human GSC cultures that was different from transcriptome analysis and better at predicting functional subgroups. In both species the chromatin accessibility-guided clusters were predominantly determined by distal regulatory element (DRE) regions, displayed contrasting sets of transcription factor binding motifs, and exhibited different functional and drug-response properties. Cross-species analysis of DRE regions in accessible chromatin revealed conserved epigenetic regulation of mouse and human GSCs. Human ATAC-seq data produced three distinct clusters with significant overlap to our previous mouse cell of origin- based stratification, where two of the clusters displayed significantly different patient survival. We conclude that epigenetic regulation of GSCs primarily is dictated by developmental origin which controls key GSC properties and affects therapeutic response.

## Introduction

Glioblastoma (GBM) is one of the most aggressive cancers and the most frequent and lethal primary malignant brain tumor ^1^. Standard therapy of care includes maximal-safe surgical resection, concomitant chemo- and radiotherapy and adjuvant chemotherapy, yet the two-year survival is 18.5% ^1^. Treatment-resistance is explained by extensive genetic and epigenetic tumor cell heterogeneity of GBM, both with regard to inter-tumor heterogeneity ^2-4^, and intra-tumor heterogeneity at different regions ^5^ and in individual cells ^6-9^. Large efforts have been done to converge GBM heterogeneity into biologically and clinically relevant subgroups of GBM. Transcriptome-based stratifications have produced three major isocitrate dehydrogenases 1 and 2 (*IDH1* and *IDH2*) wildtype (wt) GBM subtypes: proneural (PN), classical (CL), mesenchymal (MS) ^2,10,11^. Studies of patient- derived GSC cultures, clonal derivatives and single cells have shown the presence of a PN to MS differentiation axis with plasticity of the states ^8,12,13^, and a comprehensive GBM single cell analysis has uncovered additional and dynamic cellular states in GBM tumors ^7^. The GBM epigenome has been most frequently analyzed by DNA methylation profiling ^3,4,14^ and methylomes have proven prognostically more useful than transcriptomes to predict patient survival ^3,4^, demonstrating the importance of understanding of the GBM epigenome. The active chromatin landscape of GBM has been investigated with chromatin immunoprecipitation sequencing (ChIP-seq) of acetylated lysine 27 on histone H3 (H3K27ac) in a collection of primary tumors and GSC cultures ^15^, and by the assay for transposase-accessible chromatin using sequencing (ATAC-seq) ^8,16,17^, which have uncovered subgroups of GBM suggested to be regulated by different sets of transcription factors (TFs).

Several studies have implied a connection between GBM molecular subgroups and developmental origin ^2,18^. Methylation profiling has proven particularly useful to connect primary tumors with their tissue of origin ^19^ and has been used to separate GBM with higher resolution than gene expression ^4,20^. We have shown by experimental modeling of GBM that developmental state and age of the cell of origin could affect its vulnerability to GBM development ^21^, and that it shaped the phenotype of the resulting GBM stem cells (GSCs) ^21,22^. Tumors were induced by the same oncogenic events in different mouse cell lineages which produced contrasting tumor cell phenotypes with regard to malignancy and drug sensitivity, where a more differentiated origin promoted a less tumorigenic but more drug resistant mouse GSC (mGSC) phenotype ^22^. Through a cross-species GSC-based stratification approach applying the mouse cell of origin (MCO) gene signature of differentially expressed genes on a large collection of human GSC cultures, we found that developmental origin could be used to stratify functionally distinct groups of patient-derived GSC cultures ^22^. A recent similar cross-species approach has further corroborated the importance of cell lineage origin in GBM ^23^. In all this has demonstrated that inter-tumor heterogeneity to a large extent is shaped by the intrinsic properties of the GBM cell of origin which result in highly dynamic GSCs that basically evade all current therapies. To understand how cell lineage regulates these important GSC properties we performed a cross-species epigenome analysis.

We have analyzed the chromatin accessibility landscape of 9 mGSC cultures of defined developmental origin and 60 IDH wild-type human GSC (hGSC) cultures with high sensitivity ATAC-seq ^24^. We have related the results to a range of molecular and functional data and show that genome-wide chromatin accessibility is a better predictor of GSC phenotype and survival than transcriptome in both mouse and human cells. Cross-species analyses supported the subgroups to be cell lineage controlled and showed a conservation of enriched TF motifs of the differential accessible chromatin regions. Our work also highlights the value of using mouse models of different cell lineages to obtain relevant experimental coverage of GBM inter-patient heterogeneity.

## Results

### Chromatin accessibility in mouse GSCs of different origin predict self-renewal and tumorigenicity

First, we performed ATAC-seq analysis of nine previously established mGSC cultures derived from different cell lineages but induced by the same oncogenic drivers along with one NSC culture from each mouse strain ^22^ (**Figure 1a**, **Table S1**). The GSC cultures originated from mouse GBMs induced in adult tv-a transgenic mice (*G/tv-a;Arf*-/-, *N/tv-a;Arf*- /- and *C/tv-a;Arf*-/-) by RCAS-PDGFB, where the cell of origin had been deduced to be a neural stem cell (NSC)-like cell in *G/tv-a* mice, an astrocyte precursor cell (APC)-like cell in *N/tv-a* mice and an oligodendrocyte precursor cell (OPC)-like cell in *C/tv-a* mice ^22^. The ATAC-seq data from all samples was determined to be of high quality based on analyses of enrichment of sequence reads at transcription start sites (TSS), fraction of reads in peaks (FRiP) and reproducibility (**Figure 1b and Supplementary Fig. 1a, 1b)**. This was also supported by the differential chromatin accessibility at the leucine-rich repeat-containing G-protein coupled receptor 5 (*Lgr5*), Adenylyl cyclase type 2 (*Adcy2*), and 2’,3’-Cyclic-nucleotide 3’-phosphodiesterase (*Cnp*) loci (**Figure 1c**). *Lgr5* and *Adcy2* are included in the MCO gene signature ^22^, *Lgr5* as a marker of NSC-like origin cultures and *Adcy2* as marker of APC-like origin cultures. CNP expression was a prerequisite for tumor initiation in OPC-like cells of *C/tv-a;Arf-/-* mice, and although the CNP protein cannot be detected in these tumors or mGSC cultures, the pronounced open chromatin corroborated their OPC origin. Thus, the ATAC-seq profiles from all mGSC cultures accurately captured cell lineage distinctions of the mGSC cultures.

**Figure 1.**
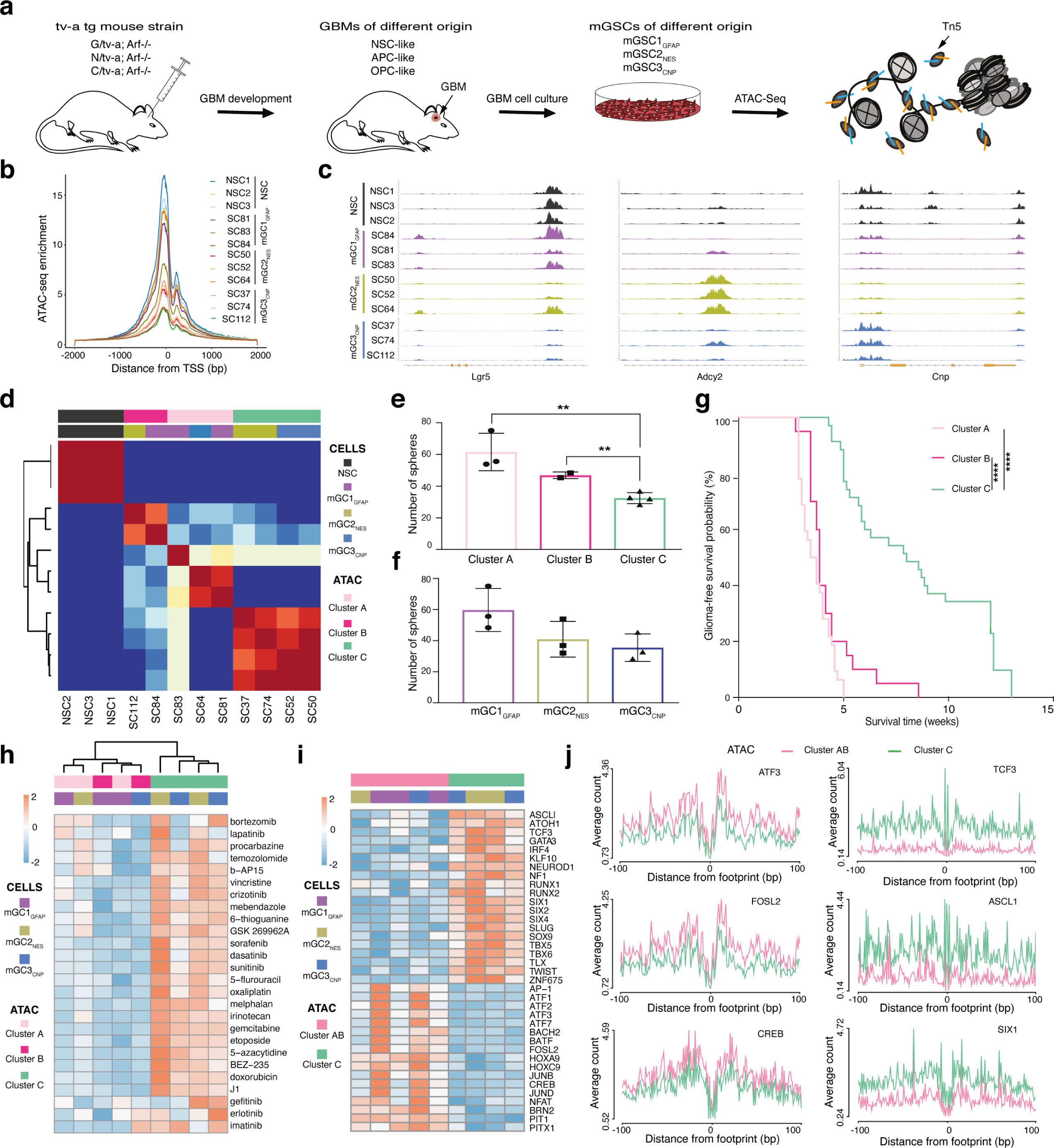
Chromatin accessibility in mouse GSCs of different origin predict self- renewal and tumorigenicity. (a) Workflow to decode chromatin accessibility in mouse GSC cultures. (b) Enrichment of ATAC-seq reads at TSS. (c) Genome browser tracks of ATAC-seq data for *Lgr5*, *Adcy2* and *Cnp* genes. (d) NMF cluster analysis of mouse ATAC-seq data. (e) Mouse GSC sphere-forming analysis comparing ATAC clusters. Student’s t-test was performed on all pair-wise comparisons, significant differences are indicated: A versus C, **p = 0.0049; B versus C, **p = 0.0066. (f) Mouse GSC sphere-forming analysis comparing cell of origin groups. Student’s t-test was performed on all pair-wise comparisons, there were no significant differences. (g) Kaplan-Meier survival analysis of glioma-free survival in syngeneic mice injected with mGSC cultures. Log-rank (Mantel-Cox) test showed significant differences between A and C, and between B and C, ****p < 0.0001 for both. (h) Heatmap of drug response scores for nine mouse GSC cultures tested for 26 anti-cancer drugs. (i) Heatmap of TF motif enrichment scores in mGSC chromatin accessible sites comparing cluster AB with C. (j) Significant TF footprints of enriched TF motifs comparing cluster AB with C. Friedman- Nemenyi test was performed, p < 0.05 for all.

Previous principal component analysis (PCA) analysis of gene expression array data from the mGSC cultures had shown a clear separation based on developmental origin ^22^. PCA analysis of ATAC-seq data did not reproduce the cell of origin groups (**Supplementary Fig. 1c**). Neither did consensus clustering (**Supplementary Fig. 1d**) nor non-negative matrix factorization (NMF) analysis (**Supplementary Fig. 1d, 1e**) of the ATAC-seq data, which made us conclude that the global chromatin accessibility within each cell of origin group had a greater variability compared with gene expression. All analyses showed a clear separation between NSCs and GSCs, and both NMF and consensus clustering divided the mGSC cultures into three clusters named A, B and C (**Figure 1d and Supplementary Fig. 1d**). The most distinctive differences between the clusters were that all NSC-like origin GSC cultures were in cluster A or B while the majority of APC-like and OPC-like origin GSC cultures were in cluster C.

To understand if the ATAC-seq clusters could be related to functional properties of the cells, we first analyzed self-renewal, a key GSC function. There were significant differences in sphere-forming ability between ATAC-seq clusters A and B compared to C, respectively (**Figure 1e**), while there were no differences for the cell of origin groups (**Figure 1f)**. We re- grouped previously generated in vivo data ^22^ using the ATAC-seq clusters and found, in line with the sphere data, significantly longer survival for mice injected with cluster C mGSC cultures compared to the other two clusters (**Figure 1g)**. Notably, all mice that were alive at the endpoint of the experiment were found in cluster C. We also re-analyzed the drug response data ^22^ and found that cluster C cultures grouped together and were overall more drug-resistant (**Figure 1h**). Taken together, we could conclude that chromatin accessibility could better predict self-renewal, tumorigenicity and drug resistance of mGSC cultures compared to the cell of origin groups. In concordance with the NMF analysis (**Figure 1d**) the functional analyses showed that cluster A and B were highly similar, so based on this we merged the ATAC-seq peaks for cluster A and B (AB) in the following analyses.

To investigate the underlying molecular regulation of the ATAC-seq clusters, first, we identified the differentially enriched ATAC peaks between cluster AB compared to cluster C (Fold change (FC) >2, false discovery rate (FDR) <0.05) (**Supplementary Fig. 1f, Table S2)**. This produced 2553 differential peaks in cluster AB, and 1232 differential peaks in cluster C. Genomic annotation showed that 87% of the differential peaks were from non-promoter regions, termed distal regulatory elements (DREs, >3 Kbp but <500 Kbp of TSS). Next, we performed TF motif enrichment analysis from the differential regions to understand the regulation of the mGSC ATAC clusters (**Figure 1i**). The dominant motifs in cluster AB were basic leucine zipper domain (bZIP) motifs, in particular of the AP-1 family (ATF1, ATF2, ATF3, ATF7, FOSL2, JUNB, JUND). AP-1 TFs play important roles in responses to extracellular stimuli, have strong connections with cancer ^25^ and have been reported to be overexpressed in patient GBM samples ^26^. In addition to bZIP motifs we also found e.g. BRN2 (encoded by *Pou3f2*) which is part of a core set of TFs essential for GBM growth ^27,28^.

In cluster C the majority of enriched TF motifs belonged to the basic helix-loop-helix (bHLH, e.g. ASCL1, ATOH1, TCF3, KLF10, NEUROD1, TWIST) or homeobox (SIX1, SIX2, SIX4) families. These are in general associated with developmental and proneural processes often promoting a more differentiated phenotype. Several studies have also shown a connection with GBM where e.g. SIX1, SOX9 and TLX, have been connected with stemness and proliferation of GBM cells ^29-31^. To further analyze TF regulation we performed TF footprint analysis of differential ATAC peaks, a computational method to predict TF binding (**Figure 1j and Supplementary Fig. 1g**). This showed a significantly higher occupancy at ATF3, FOSL2 and CREB motifs in cluster AB (**Figure 1j**), and significantly higher occupancy of ASCL1, TCF3, SIX1, SIX2, SIX4, SLUG and TWIST motifs in cluster C (**Figure 1j and Supplementary Fig. 1g**). Other TF motifs that were clearly different (non-significant) between the groups were BRN2 and NEUROD1 (**Supplementary Fig. 1g**). These analyses provided a molecular basis for the functionally different mGSC ATAC clusters and suggested the presence of different transcriptional regulation.

Lastly, to characterize the mGSC ATAC clusters we used the available gene expression array data ^22^ and performed gene set enrichment analysis (GSEA) to compare cluster AB to cluster C. We found that the “Wong embryonic stem cell core” gene set, showed enrichment for cluster AB and inverse enrichment for cluster C (**Supplementary Fig. 1h**), in agreement with the different self-renewal capacities of the cultures. There was no difference for the “Beier glioma stem cell up” gene set (**Supplementary Fig. 1i**), consistent with the cancer stem cell phenotype of all mGSC cultures.

In summary we found that chromatin accessibility profiling is a more accurate molecular method to detect heterogeneity among mGSC cultures of the same origin compared to transcriptome profiling. This heterogeneity could also more precisely predict self-renewal, malignancy and drug response capacities of the cultures, which showed the importance of cell lineage-regulation of GSCs. The fact that a large proportion of the differential chromatin accessible regions were DREs enriched with distinct sets of TF motifs provided additional support for discrete regulation of the mGSC clusters.

### Heterogeneous chromatin accessibility across 60 patient-derived GSC cultures

Next, we investigated the chromatin accessibility landscape in our local human GBM cell culture (HGCC) biobank ^22,32,33^. We performed ATAC-seq on 60 patient-derived IDH wildtype GSC cultures (hGSC) (**Figure 2a**, **Table S1**). Again, we applied established and stringent criteria for the ATAC-seq data processing and could show high Pearson correlation coefficient (0.8-0.98) for technical replicates, TSS enrichment scores above 3.8, and a FRiP of all 60 samples above 10% ^34^ (**Supplementary Fig. 2a-c**). We performed a saturation analysis using random sampling non-linear regression to analyze the fraction of all predicted accessible chromatin regions that we could expect to detect with 60 cultures (**Figure 2b**).

**Figure 2.**
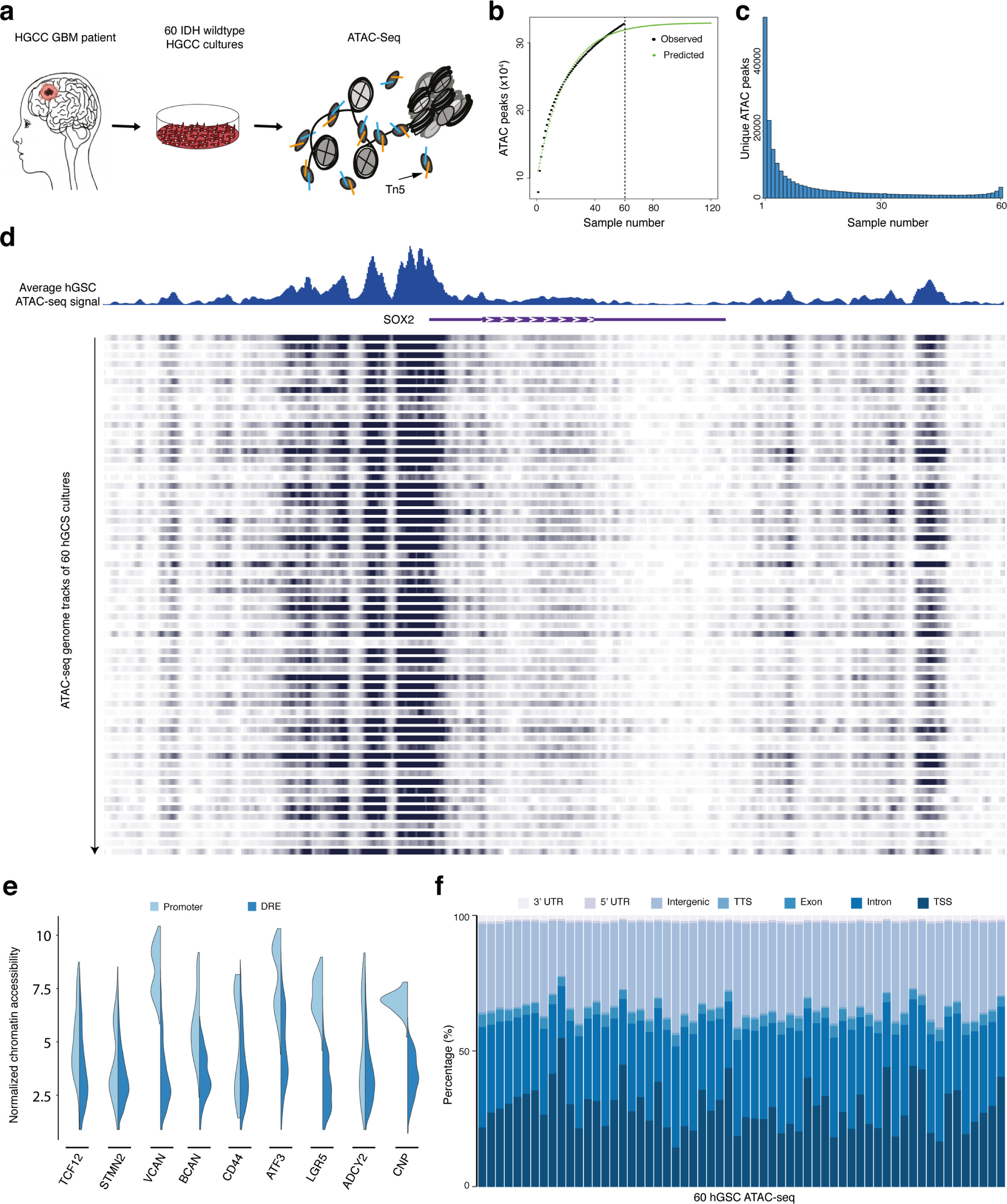
Heterogeneous chromatin accessibility across 60 patient-derived GSC cultures. (a) Workflow to decode the chromatin accessibility of 60 patient-derived IDH wildtype GSC cultures. (b) Saturation analysis using a non-linear model. Total number of predicted accessible chromatin regions in GBM (light green line), observed number of accessible chromatin regions in 60 hGSC samples (dark green line). (c) Histogram of the distribution of unique ATAC peaks in the cohort. (d) Genome browser tracks of ATAC-seq signal at *SOX2*. Top panel shows the average ATAC-seq signal of all 60 samples. Bottom panel shows the ATAC-seq signal from each individual hGSC culture. (e) Violin plots of cohort-wide distribution of chromatin accessibility at promoters and DRE regions of GBM meta module and cell lineage-relevant genes. (f) Genomic annotation of ATAC peaks in each hGSC sample.

This showed that our sample size was large enough and that all predicted regions of accessible chromatin would be detected with 47 cultures. In total, we had captured 323526 ATAC peaks from our hGSC cohort. To obtain an overview of the global chromatin accessibility landscape of the samples we identified all unique chromatin-accessible regions in the entire data and calculated for each region the number of samples it was present in (**Figure 2c**). A large proportion (25.4%) of ATAC peaks was only detected in one hGSC culture, and just 1.5% of the accessible regions were common to all 60 cultures.

Nonetheless, unique chromatin-regions showed increased openness the more frequently they were present in the samples (**Supplementary Fig. 2d**). *SOX2* showed an overall high chromatin accessibility across the cohort (**Figure 2d**) in line with previous data showing SOX2 expression in all HGCC cultures investigated ^35^. Yet, individual cultures showed a clear variability in chromatin openness of this loci (**Figure 2d**). Inter-culture heterogeneity was further sustained by analyses of promoter (-1 Kbp to +100 bp of TSS) and DRE regions of GSC meta module genes ^7^ and MCO genes ^22^ (**Figure 2e and Supplementary Fig. 2e**). Also structural genomic annotation of the ATAC data showed a clear variation in chromatin- openness among the 60 cultures (**Figure 2f**). Taken together, this demonstrated that our cohort of 60 patient-derived GSC cultures displayed a highly heterogeneous chromatin accessibility landscape.

### Chromatin-accessibility robustly identify three clusters of patient-derived GSC cultures

To identify unifying features of the hGSC cohort we performed NMF analysis on the ATAC-seq data, which produced three clusters: ATAC60-C1 (n=22), ATAC60-C2 (n=16) and ATAC60-C3 (n=22) (**Supplementary Fig. 3a, 3b, Table S1**). A large part of the HGCC cultures had previously been classified based on gene expression according to The Cancer Genome Altlas (TCGA) subtypes ^35^ and by using the MCO gene signature (MCO 1-3) ^22^. To compare the ATAC-seq clusters with the TCGA and MCO classifications we excluded hGSC samples lacking such information and re-analyzed 50 samples with NMF. This produced, again, three clusters: ATAC50 C1 (n=19), ATAC50 C2 (n=14) and ATAC50 C3 (n=17) (**Figure 3a and Supplementary Fig. 3c**). Comparing ATAC50 to ATAC60 clusters showed that only three samples had changed cluster in ATAC50 (**Supplementary Fig. 3d, Table S1**), which indicated a robustness of the chromatin accessibility-based clustering. From hereon we only refer to the ATAC50 classification.

**Figure 3.**
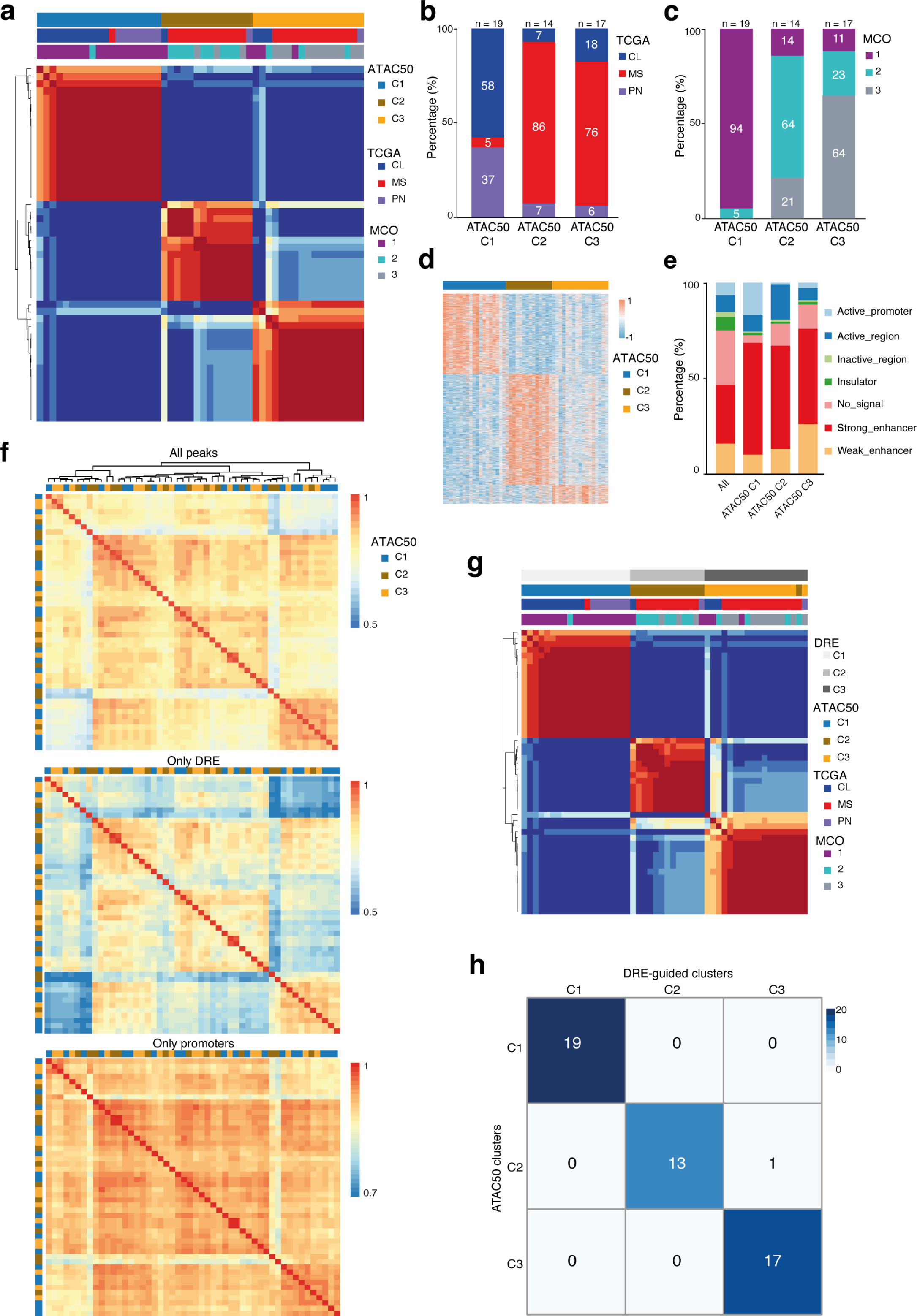
Chromatin-accessibility robustly identify three clusters of patient-derived GSC cultures. (a) NMF cluster analysis of ATAC-seq data from 50 patient-derived GSC cultures. (b) Distribution of TCGA subtypes among ATAC50 clusters. (c) Distribution of MCO subgroups among ATAC50 clusters. (d) Heatmap of unique ATAC peaks of each ATAC50 cluster. (e) Genomic annotation of all ATAC peaks in the hGSC cohort and unique ATAC peaks of each ATAC50 cluster. (f) Pearson correlation heatmaps of all ATAC peaks (top), ATAC peaks of DREs (middle) and ATAC peaks of promoters (bottom). Samples are arranged in the same order in all heatmaps, based on Pearson correlation of all ATAC peaks. (g) NMF clustering of DRE ATAC peaks from 50 human GSC cultures. (h) Overlap of ATAC50 clusters with DRE-guided clusters.

When comparing the ATAC50 clusters with the TCGA subtypes there was little overlap (**Figure 3b**). The majority of PN and CL cultures were in C1 while MS cultures basically were divided between C2 and C3. Comparing to MCO showed a higher degree of overlap (**Figure 3c**), likely reflecting the relation between developmental origin and epigenetic state of GSC.

Next, we extracted the unique chromatin-accessible regions for each ATAC50 cluster with DESeq2 (FC >2, FDR <0.001, peak average intensity >30, and coefficient of variance <0.2) (**Figure 3d**). By this we identified 4023 regions in C1, 5547 in C2 and 949 in C3 (**Figure 3d**, **Table S3**). The genomic features of all chromatin-accessible regions from each ATAC50 cluster were annotated using the chromatin state discovery and characterization software (ChromHMM) ^36^ (**Figures 3E and Supplementary Fig. 3e**). There was a diverse distribution of chromatin states with some marked differences between ATAC50 clusters. C1 had the largest proportion of active promoter regions (H3K4me3 and H3K27ac), C2 occupied a higher proportion of active regions (H3K27ac), and C3 had a higher frequency of weak enhancer regions (H3K4me1). Common to all three clusters was that the combined proportion of strong and weak enhancer regions constituted the biggest proportion of chromatin states, clearly larger compared to the distribution in the whole ATAC-seq data (**Figure 3e**). This indicated, as in mGSCs, that DRE regions were central in defining the ATAC50 clusters. To test our hypothesis, we performed Pearson correlation hierarchal clustering of all ATAC peaks, of DRE regions only, and of promoter regions only (**Figure 3f**). All ATAC peaks and DRE peaks displayed similar dynamic range of chromatin openness and cluster patterns, while the dynamic range of promoter regions was smaller which supported our assumption. NMF clustering of DRE ATAC peaks (**Figure 3g and Supplementary Fig. 3f**) produced almost identical clusters as ATAC50 (**Figure 3h**).

Clustering promoter regions resulted in two clusters (**Supplementary Fig. 3g, 3h**) that were entirely different and non-overlapping to ATAC50 (**Supplementary Fig. 3i**). Collectively, our analyses showed that chromatin accessibility could robustly stratify hGSC cultures and clusters were predominantly dictated by the DRE regions. The high correspondence of the ATAC50 clusters with the MCO stratification implied an important role of cell lineage-controlled gene regulation of human GSC cultures.

### ATAC50 clusters are phenotypically distinct

To phenotypically characterize the ATAC50 clusters we first used hGSC gene expression array data ^22^ and analyzed the 256 GSC meta module genes ^7^ across the 50 cultures (**Figure 4a and Supplementary Fig. 4**). While only 53 genes showed a significant difference between the clusters (**Figure 4b**) there were clear differences comparing global meta module gene expression (**Figure 4a**). C1 showed significantly higher expression of NPC1, NPC2 and OPC genes compared to both C2 and C3, significantly higher expression of AC genes compared to C2, and significantly lower expression of MES genes compared to both C2 and C3. Thus, C1 and C2 were always at the end of the spectrum with C3 in the middle. Notably, C3 showed significantly higher expression of NPC1, OPC and AC genes and significantly lower expression of MES2 genes compared to C2. This suggested that ATAC50 clusters were separated along a gradient of GSC states with C1 being progenitor cell-like, C2 being mesenchymal-like and C3 being intermediate.

**Figure 4.**
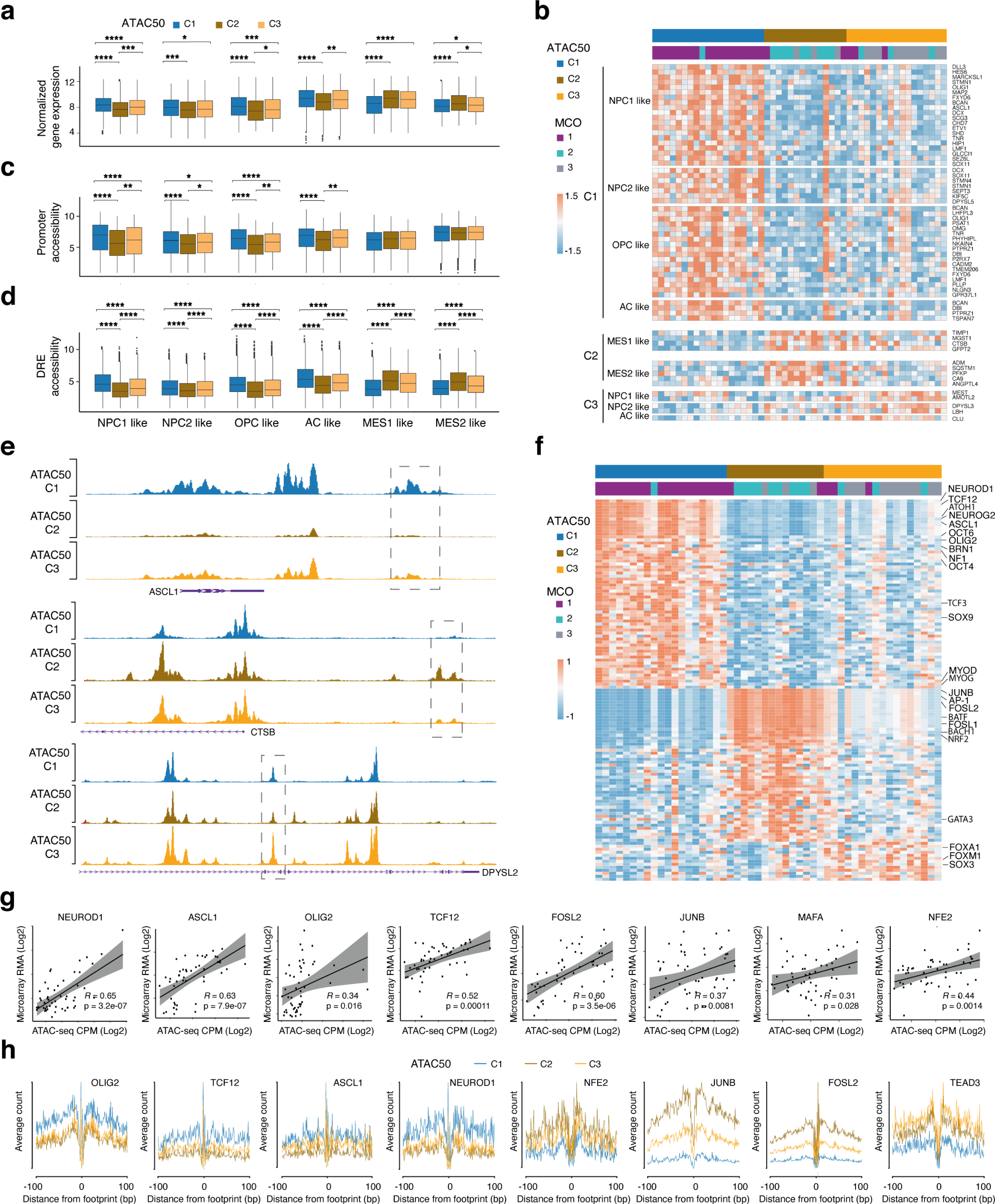
hGSC ATAC50 clusters are phenotypically distinct. (a) Average gene expression of each GSC meta module comparing ATAC50 clusters. (b) Gene expression of ATAC50 cluster significant meta module genes. (c) Chromatin accessibility at promoters of GSC meta module genes. (d) Chromatin accessibility at DREs of GSC meta module genes. Student’s t-test was performed on all pair-wise comparisons in (**a**), (**c**) and (**d**), significant differences are indicated, *p < 0.05, **p < 0.01, ***p < 0.001, ****p < 0.0001. (e) Genome tracks for *OLIG2*, *NDRG1*, *LBH*. (f) Heatmap of significantly enriched cluster specific TF motifs. (g) Scatter plots of TF motif chromatin accessibility (x-axis, normalized chromatin openness) and TF gene expression (y-axis, microarray data, counts per million reads mapped (CPM)) of significantly enriched TF motifs in (**f**). *R* = Pearson correlation, p calculated by Student’s t- test. (h) TF footprint analysis of cluster-specific ATAC50 peaks. Friedman-Nemenyi test was performed, p < 0.05 for all.

Next, we analyzed chromatin openness of meta module genes in promoter regions (**Figure 4c**) and DRE regions (**Figure 4d**). The openness of DRE regions was significantly different between all clusters in all meta modules whereas promoter regions showed less distinct differences for NPC1, NPC2, OPC and AC meta modules and were non-significant for MES1 and MES2. This is in line with the dominant role of DRE regions to separate the ATAC50 clusters (**Figure 3f**).

Of the meta module genes with significant different gene expression (**Figure 4b**) the majority (39) were higher expressed in C1 and belonged to the NPC1, NPC2, OPC and AC modules.

C2 showed a higher expression of some MES1 and MES2 genes, and C3 cultures displayed higher expression of a few NPC1, NPC2 and AC genes. To investigate the chromatin openness of DRE regions of these genes we first linked all human ATAC DRE regions with their nearest gene through a peak-to-gene linking prediction analysis ^34^ (**Supplementary Fig. 5a, Table S4**). This was used to compare ATAC peaks of the significant genes (**Figure 4b**) between clusters (**Table S4**) which showed that 47% (25 of 53) had significantly different chromatin accessibility in DRE regions (**Figure 4e**, **Table S4**), compared to only 7 of 53 for promoter regions, supporting the importance of DREs in regulating ATAC50 clusters.

To investigate underlying mechanisms controlling the ATAC50 clusters, we performed TF motif enrichment analysis on significantly different chromatin-accessible regions. Of the top- 50 most variable TFs motifs the majority were bZIP (n=15) or bHLH (n=15) motifs (**Supplementary Fig. 5b, Table S5**). There was a clear and inverse enrichment when comparing ATAC50 C1 and C2, with bHLH motifs being most common in C1 and bZIP motifs being most common in C2. GSCs maintained in stem cell media have been shown to enrich for bHLH TFs while serum media enriched for bZIP TFs ^27^ corroborating different state identities of C1 and C2 cultures. To find distinctive features of each cluster we extracted the significantly enriched cluster-specific TF motifs (**Figure 4f**, **Table S6**). This identified 64 uniquely enriched motifs in C1, 51 in C2, and 13 in C3. Among the TFs in C1 many were regulators of neural development with strong connections to GBM such as TCF12 ^37-40^, ASCL1 ^41^, OLIG2 ^42^ and SOX9 ^43^. Dominant TF motifs enriched in C2 were AP- 1 complex motifs of the JUN, FOS, ATF and MAF families and motifs of the MAF dimerizing proteins NRF2, BACH1 and BACH2, which have also been associated with cancer progression and metastasis ^25^ and are plausible candidates to regulate the mesenchymal features of C2 cultures. Since C3 was intermediate to C1 and C2, there were fewer uniquely enriched TF motifs in this cluster. Among them were GBM-associated FOXM1, FOXA1 and SOX3 (**Figure 4f**) ^37-40,44^. When analyzing the relationship of TF motif openness and TF gene expression we found a positive correlation for several TFs (**Figure 4g**) which strengthened their involvement in shaping the cluster phenotypes. We also analyzed TF occupancy by footprint analysis of differential peaks (**Figure 4h and Supplementary Fig. 5c**). We found that ATAC50 C1 showed significantly higher occupancy of OLIG2, TCF12, ASCL1 and NEUROD1 compared to C2 and C3, while C2 and C3 showed significantly higher occupancy for NF-E2, JUNB and FOSL2 compared to C1. Among the C3-uniquely enriched TF motifs there were no significant footprints, but among the top-50 variable TF motifs TEAD3 showed significantly higher occupancy for C3 compared to C1 (**Figure 4h**). In all, the TF motif enrichment, TF gene expression and TF occupancy analyses confirmed the phenotypic differences and revealed distinct epigenetic regulation of the ATAC50 clusters.

### ATAC50 classification produce functional separation of hGSC cultures

Since epigenetic clusters of mGSCs efficiently distinguished functional groups, we asked whether ATAC50 clusters also would. We investigated essential GSC properties in 16 C1, 8 C2 and 13 C3 cultures, of which the majority had overlapping MCO and ATAC50 classifications (**Table S1**). We first performed consecutive sphere-forming assays under clonal conditions (**Figure 5a**). C1 cultures displayed the highest sphere-forming ability while there was no significant difference between C2 and C3 cultures, although C3 cultures produced a higher average number of spheres. Extreme limiting dilution assay (ELDA) showed that C1 cultures had the highest self-renewal capacity but there was also a difference between C2 and C3 (**Figure 5b**). A similar result was found for cell proliferation where C1 had a significantly higher BrdU incorporation compared to both C2 and C3 cultures, with C3 showing intermediate proliferation (**Figure 5c**). Migration was analyzed with the spheroid collagen gel invasion assay. Here we found that C2 cultures were significantly more invasive than C1 and C3 cultures (**Figure 5d**). In all, these functional characteristics were in accordance with the stem and progenitor cell-like molecular phenotype of C1 cultures, the mesenchymal-like phenotype of C2 culture and the mixed molecular phenotype of the C3 cultures.

**Figure 5.**
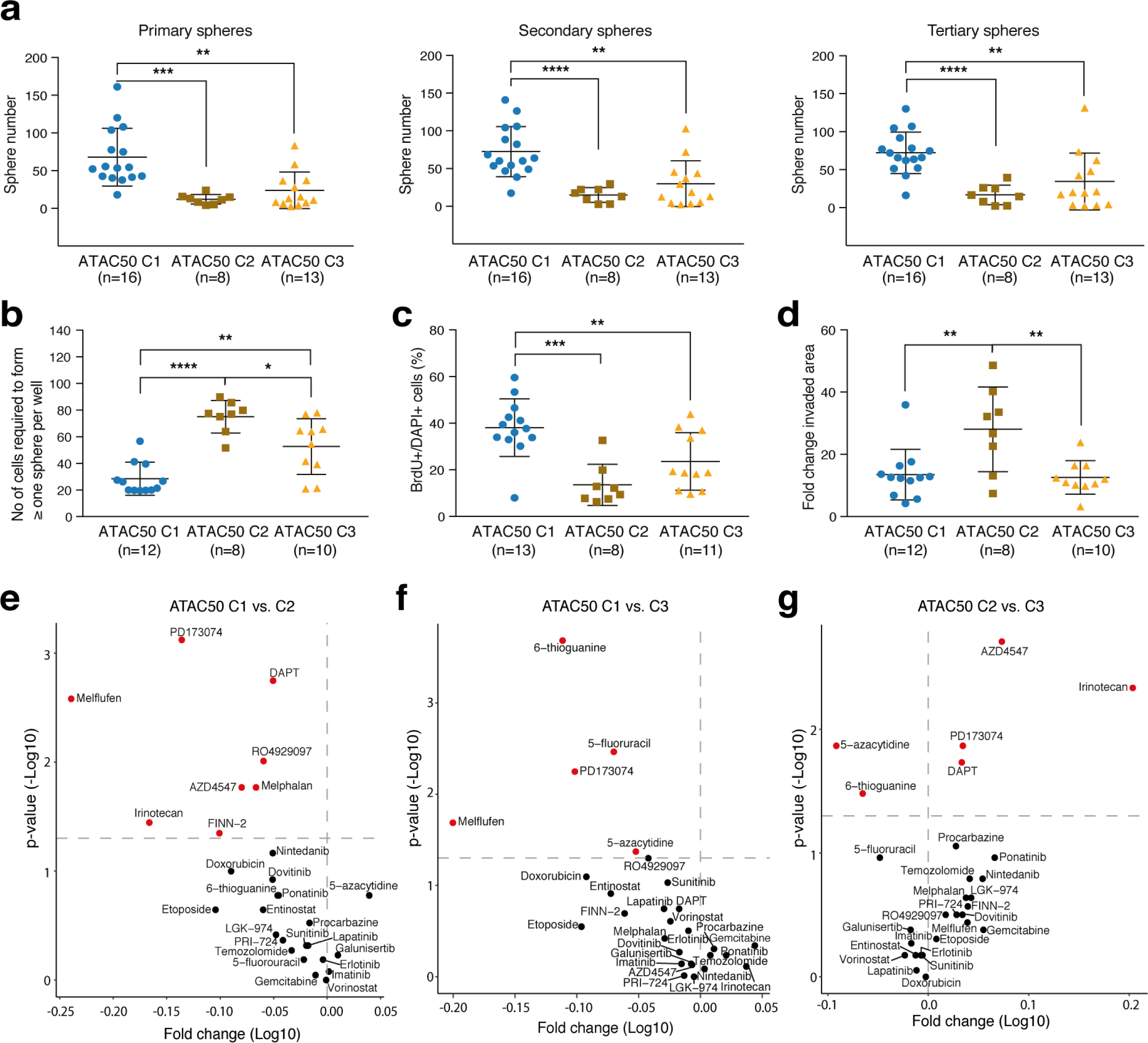
ATAC50 classification produce functional separation of hGSC cultures. (a) Sphere assays comparing ATAC50 clusters. Data show mean ± SEM. Student’s t-test was performed on all pair-wise comparisons, significant differences are indicated. Primary spheres: C1 versus C2, ***p = 0.0005; C1 versus C3, **p = 0.0013. Secondary spheres: C1 versus C2, ****p < 0.0001; C1 versus C3, **p = 0.0014. Tertiary spheres: C1 versus C2, ****p < 0.0001; C1 versus C3, **p = 0.0041. n, number of different cell cultures analyzed. (b) ELDA comparing ATAC50 clusters. Data show mean ± SEM. Student’s t-test: C1 versus C2, ****p < 0.0001; C1 versus C3, **p = 0.0021; C2 versus C3, *p = 0.0169. n, number of different cell cultures analyzed. (c) Frequency of BrdU positive cells. Data show mean ± SEM. Student’s t-test was performed on all pair-wise comparisons, significant differences are indicated: C1 versus C2, ***p = 0.0001; C1 versus C3, **p = 0.0088. n, number of different cell cultures analyzed. (d) Collagen invasion assay measuring the invaded area at 24 hours. Data show mean ± SEM. Student’s t-test was performed on all pair-wise comparisons, significant differences are indicated: C1 versus C2, **p = 0.0072; C2 versus C3, **p = 0.0043. n, number of different cell cultures analyzed. (**e-g**) Volcano plots of pair-wise comparisons of AUC scores in ATAC50 C1 (n=12), C2 (n=7) and C3 (n=10) cultures to 28 anti-cancer drugs. Red circles indicate significantly different drug response. n, number of different cell cultures analyzed. (e) ATAC50 C1 versus C2. Red circles in upper left corner, C1 is more sensitive. (f) ATAC50 C1 versus C3. Red circles in upper left corner, C1 is more sensitive. (g) ATAC50 C2 versus C3. Red circles in upper left corner, C2 is more sensitive. Red circles in upper right corner, C3 is more sensitive.

We also analyzed the drug response phenotype of 11 C1, 7 C2 and 10 C3 cultures by measuring cell viability after 72 hours exposure to a collection of 28 anti-cancer drugs at 7 different concentrations (**Figure 5e-g, Table S7**). This produced dose-response curves that were converted to area under the curve (AUC) measures that were compared pair-wise between clusters. There was a clear overall higher sensitivity of C1 cultures to the compounds compared to both C2 (**Figure 5e**) and C3 (**Figure 5f**) cultures. All drugs that produced a significantly different response between C1 and C2 or C3 cultures were more effective in C1 cultures. These comparisons identified two compounds as particularly efficient for C1 cultures, Melflufen (alkylating) and PD173074 (FGFR1 inhibitor). When comparing C2 to C3 cultures, C3 cultures were clearly, overall, more sensitive to the tested drugs (**Figure 5g**). However, C3 cultures showed a significantly higher resistance to two drugs, 5-azacytidine and 6-thioguanine, compared to both C1 and C2 cultures (**Figure 5f and 5g**). The distinct drug response phenotypes of the ATAC50 clusters suggested that cell lineage dependencies are important to account for when developing therapeutic strategies for GBM.

### ATAC50 clusters exhibit different mouse and patient survival

Orthotopic tumor growth is a defining capacity of cancer stem cells. We used in vivo data, in total 322 intracranially injected immune-deficient mice, from published ^22,32,33,35^ and unpublished experiments, and included only individuals that had been killed because of disease symptoms before the experimental endpoint (**Table S8**). When grouping mice according to the ATAC50 clusters we found a significant difference in survival between all groups with C1 being most aggressive (**Figure 6a**). We also analyzed survival of GBM patients from whom hGSC cultures had been derived (**Figure 6b-d**). We first grouped patients according to the widely used TCGA classification (**Figure 6b**), which showed no difference between the groups. Nor did the ATAC50 clusters (**Figure 6c**), although the curves seemed slightly more separated. We also analyzed the MCO clusters because of the high degree of overlap with ATAC50 (**Figure 6d**). This showed a significant survival difference between MCO2 and MCO3 patients. When we combined the MCO and ATAC50 classifications, i.e. included only those patients whom showed an overlap for both, the significant survival benefit of C3 patients compared to C2 remained (**Figure 6e**). The differences between mouse and patient survival can have many explanations, e.g. clonal selection of GSCs, but we believe that one important factor was the effect of therapy in the patients. Although there were no significant differences between clusters when comparing drug-response scores for temozolomide we noted that C3 cultures were always most sensitive to this drug compared to C1 and C2 (**Figure 5e-g**). When we analyzed the dose-response curves for temozolomide we found that C2 cultures were most resistant and C3 cultures most sensitive at all concentrations, with significant difference between C2 and C3 at the four highest doses (**Figure 6f**). Thus, the elevated tumorigenic properties of C1 GSCs may in patients have been counteracted by their higher sensitivity to temozolomide, and vice versa for C2 GSCs. The significantly improved survival of C3 patients could be explained by C3 cultures being least malignant (**Figure 5a-d and Figure 6a**) and most temozolomide-sensitive (**Figure 6f**). This showed that epigenetic and cell lineage-based classifications could be valuable to predict GBM patient survival, likely because of their ability distinguish key tumor cell phenotypes.

**Figure 6.**
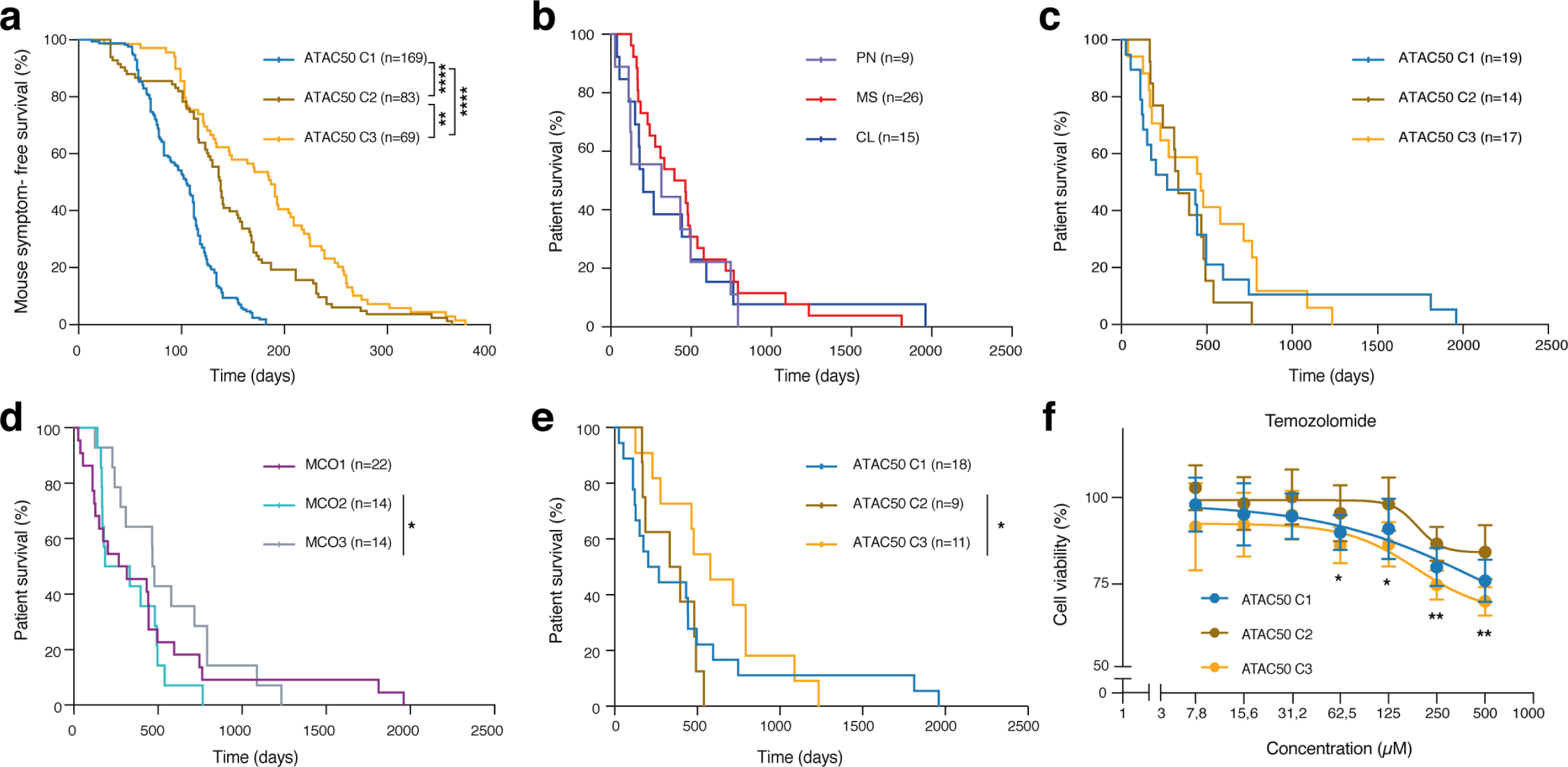
ATAC50 clusters exhibit different mouse and patient survival. (a) Kaplan-Meier analysis of symptom-free survival of intracranially injected immune- deficient mice (14 C1 cultures, 11 C2 cultures, 9 C3 cultures). Log-rank (Mantel-Cox) test: C1 versus C2 or C3, ****p < 0.0001; C2 versus C3, **p = 0.0062. n, number of injected mice. (**b-e**) Kaplan-Meier analysis of patient survival. Log-rank (Mantel-Cox) test was performed, significant differences are indicated. n, number of patients. (b) Patients were divided based on TCGA subtype. (c) Patients were divided based on ATAC50 clusters. (d) Patients were divided based on MCO classification. C2 versus C3, *p = 0.0182. (e) Patients with overlapping ATAC50 and MCO classifications. C2 versus C3, *p = 0.0371. (f) Dose-response curves show cell viability after 72 hours exposure to temozolomide. Student’s t-test was performed on all pair-wise comparisons at all concentrations, significant differences between C2 and C3 are indicated: 62,5μM, *p = 0.0405; 125 μM, *p = 0.0170; 250μM, **p = 0.0012; 500μM, **p = 0.0027.

### Cross-species analyses reveal mouse cell of origin prediction of hGSC ATAC clusters

At last, we performed cross-species analyses of chromatin accessibility in mGSC and hGSC cultures. The high overlap of the ATAC50 clusters with the cell lineage-based MCO stratification (**Figure 3c and Figure 7a**) suggested an important role of developmental regulation. We started by investigating if the chromatin landscape of the MCO genes could guide the ATAC50 clusters. Of the 196 MCO genes ^22^ we used 166 human homologues for which the ATAC-seq peaks of promoter regions were extracted and analyzed by NMF (**Supplementary Fig. 6a, 6b)**. This produced a poor overlap with the ATAC50 clusters (**Figure 7b**), consistent with the importance of DRE regions (**Figure 3h**). Next we investigated the DRE regions of the MCO genes through the peak-to-gene linking prediction analysis (**Table S4**). The MCO human homologue genes were annotated to 786 ATAC peaks that were analyzed by NMF (**Supplementary Fig. 6c, 6d**), which showed a higher concordance with ATAC50 (**Figure 7c**) compared to promoter-guided clusters (**Figure 7b**) but lower than the MCO stratification (**Figure 7a**). Finally, we used cluster-specific mouse ATAC peaks (n=3785, **Table S2**) that were annotated to 2456 mouse genes, converted to 2128 human homologue genes of which 1298 were present in the peaks-to-genes data and could be linked to 8187 human ATAC peaks. The DREs of these peaks were used in NMF (**Supplementary Fig. 6e, 6f**) which produced clusters that, surprisingly, showed a very high agreement with the ATAC50 clusters (**Figure 7d**). As reference we compared this to 1000 NMF analyses of 8187 randomly selected ATAC peaks from the peaks-to-genes data which did not reproduce the ATAC50 clusters (**Supplementary Fig. 6g**). To investigate the basis for the overlap of mouse-dictated human ATAC50 clusters, we compared enriched TF motifs in the 3785 mouse ATAC peaks to those of the corresponding 8187 human ATAC peaks.

**Figure 7.**
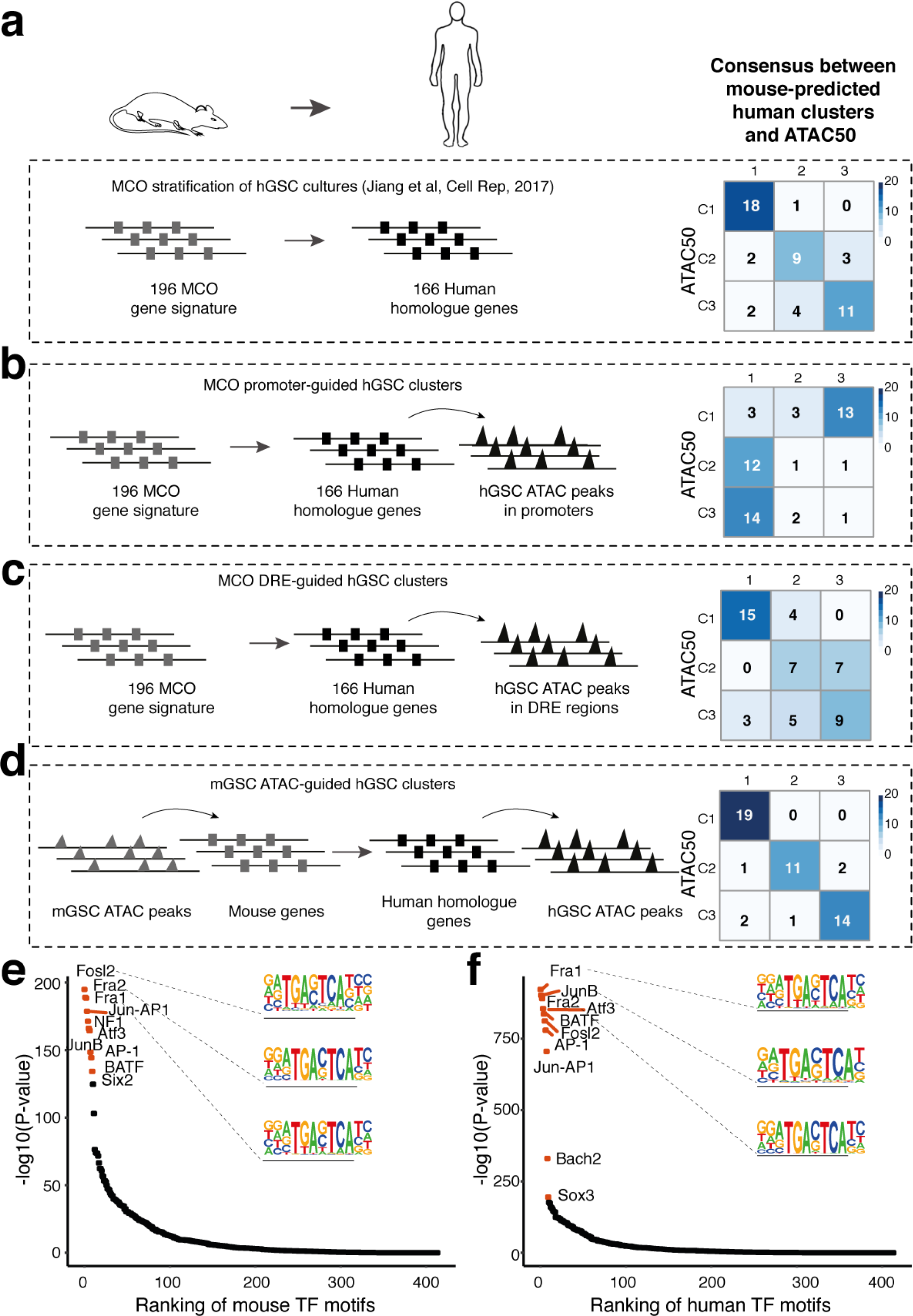
Cross-species analyses reveal conservation between mouse and human GSC chromatin accessibility. (**a-d**) Analyses of mouse GSC-guided clustering of human GSCs and their overlap with ATAC50 clusters. (a) Overlap with MCO stratification (Jiang et al, Cell Rep, 2017). (b) Overlap with MCO promoter-guided ATAC clusters. (c) Overlap with MCO DRE-guided ATAC clusters. (d) Overlap with mGSC cluster-unique ATAC-guided clusters. (e) Ranking of enriched TF motifs in mGSC cluster-specific ATAC peaks. Red circles indicate top-10 significant motifs. (f) Ranking of enriched TF motifs in the mGSC ATAC-guided hGSC ATAC peaks. Red circles indicate top-10 significant motifs.

This showed an 80% overlap in the top-10 significantly enriched TF motifs in mouse and human ATAC data (**Figure 7e, 7f**), and all overlapping motifs were of the AP-1 family. As a reference, we compared this to a 1000 times repeated control experiment where enriched TF motifs in 3785 randomly selected mouse ATAC peaks and their corresponding human ATAC peaks were compared (**Supplementary Fig. 6h**). The TF motif overlap extended from 0 to 6 with an average overlap of 22.3% (standard deviation =13.4%) and overlapping TF motifs were not restricted to the AP-1 family.

In summary, this showed that the chromatin accessibility based cross-species analysis with high precision could predict the ATAC50 clusters, with complete accuracy for C1 and with slight deviations for C2 and C3, consistent with the more similar phenotypes of C2 and C3. TFs motif analysis displayed common TFs regulatory mechanisms of mouse and human GSCs, which provide strong support for the presence of conserved cell of origin-determined epigenetic regulation of GBM.

## Discussion

The contribution of developmental regulation in GBM biology and GSC function remains to a large extent to be deciphered. We have addressed it by performing genome-wide analysis of chromatin-accessibility in mouse and human GSC cultures. Mouse GSC cultures were derived from GBMs of different cell lineages ^22^, and within each cell lineage group we found a variability in chromatin openness which we had not observed with gene expression. One explanation could be that it reflected a differentiation state variability of the targeted cell of origin since chromatin accessibility have been shown to precede changes in gene expression ^45^. Strikingly, the chromatin accessibility variability was shown to better determine essential GSC functions.

The ATAC-seq data divided both mouse and human cells in molecularly and functionally distinct groups, and these groups were mainly determined by the chromatin state of DREs and regulated by contrasting sets of TFs. Our finding of three hGSC ATAC clusters aligned with recent ATAC-seq analyses of human GBM cells and GSC cultures from independent cohorts ^8,16^ supporting our non-linear regression analysis which predicted that our cohort would be large enough to capture the spectrum of GBM inter-patient heterogeneity.

Expression of GSC state markers ^7^ and analyses of TF circuits showed that C1 cultures were glial stem and progenitor cell-like, C2 profoundly mesenchymal-like and C3 had an intermediate phenotype with mostly astrocytic and mesenchymal traits. ATAC50 clusters were also functionally well-defined where C1 cultures were most self-renewing, proliferative and tumorigenic, C2 cultures most invasive and temozolomide-resistant, and C3 cultures least tumorigenic and temozolomide-resistant. The precision of chromatin accessibility to separate TCGA mesenchymal subtype cultures into two groups with significantly different survival emphasized the importance of understanding the epigenetic regulation of the ATAC50 clusters.

The considerable overlap of the MCO classification with the ATAC50 clusters implied a cell lineage controlled regulation of hGSCs. This was corroborated by the cross-species analysis where chromatin accessibility of cluster-specific DRE regions from mGSCs could almost completely predict the ATAC50 clusters. The 80% overlap of top-10 enriched TF motifs in the cross-species comparison of accessible chromatin regions provided strong support for a conserved epigenetic cell lineage regulation of GBM. Our cross-species analysis also validated the PDGF-driven mouse GBM models as highly relevant, representing the breadth of developmental regulation present in our collection of 50 patient-derived GSC cultures. The fact that one oncogenic driver (PDGFRA activation) could reproduce the epigenetic heterogeneity of human GBM was in line with results from the comprehensive single cell RNA-seq analysis of human GBM ^7^ where multiple cellular states were shown to be present in all investigated tumors, while state distributions were proposed to be dictated by certain genetic factors such as PDGFRA. Taken together, this would argue for that GBM epigenetic heterogeneity is mainly be the consequence cell of origin-inherited developmental regulation which in turn provide the basis for possible GSC states, and that GBM driver mutations determine the state transition dynamics.

We show the power of a chromatin accessibility-based functional classification of GSCs. Continued work to identify the key regulatory elements in the DREs dictating the different properties and common features of the epigenetic clusters, and to validate key TF circuits regulating GSC states by perturbation strategies will be crucial to pinpoint therapeutic targets. Our analysis of chromatin accessibility in mGSCs and hGSCs has revealed a species conservation of the GBM epigenome and demonstrated the importance of cell lineage diversity for accurate in vivo modeling of inter-patient heterogeneity.

## Author contributions

L.U. and X.C. conceptualized, designed and supervised the study. N.P.M. planned, performed and analyzed most experiments. M.J., L.Z., Y.X., E-J.T. and X.C. performed additional experiments. X.L. performed all computational analyses with help from P.X. M.F. provided technical and material support. L.U. and X.C. wrote the manuscript with input from all co-authors.

## Acknowledgements

We thank members of the Chen and Uhrbom laboratories for scientific discussions. This work was supported by the Swedish Research Council (2016-06794, 2017-02074 to X.C., 2018-02906 to L.U.), the Swedish Cancer Society (15 0877, 18 0763 to L.U.), Beijer Foundation (to X.C.), Jeansson’s Foundation (to X.C.), Petrus och Augusta Hedlunds Stiftelse (to X.C.), Göran Gustafsson’s prize to younger researchers (to X.C.), Vleugel Foundation (to X.C.), and Uppsala University (to X.C.).

## Availability of the data

The accession number for mGSC and hGSC ATAC sequencing data generated for this study is GSE163853.

## Methods

### ATAC-seq

Supplementary Figures 1-6 are in the separated files. Table S1-8 are in the separated file.

Human GSCs, mouse GSCs and mouse NSCs were fixed with 1% formaldehyde (Thermo Fisher scientific, 28906) for 10 min and quenched with 0.125 M glycine for 5 min at room temperature. After the fixation, ATAC-seq was performed as previous described^1^. Cells were counted and 50 000 cells were used per ATAC-seq reaction. The transposition reaction followed the normal ATAC-seq protocol. After transposition, a reverse crosslink solution (final concentration 50mM Tris-Cl PH 8.0 (Invitrogen, 15568-025), 1mM EDTA (Invitrogen, AM9290G), 1% SDS (Invitrogen, 15553-035), 0.2M NaCl (Invitrogen, AM9759) and 5ng/ul proteinase K (Thermo Scientific, EO0491)) was added up to 200 µl. The mixture was incubated at 65 °C with 1200 rpm shaking in a heat block overnight, then purified with MinElute PCR Purification kit (QIAGEN, 28004) and eluted in 10 µl Qiagen EB elution buffer. Sequencing libraries were prepared following the original ATAC-seq protocol^2^. The sequencing was performed on Illumina NovaSeq 6000, and at least 20 million paired-end sequencing reads were generated for each ATAC-seq library.

### ATAC-seq data processing

After the Adapter sequence trimming, the ATAC-seq sequencing reads were mapped to genome hg19 or mm9 using bowtie2^3^. Mapped paired reads were corrected for the Tn5 cleavage position with shifting +4/-5 bp depending on the strand of reads. All mapped reads were extended to 50 bp centered by Tn5 offset. The PCR duplication were removed using Picard (http://broadinstitute.github.io/picard/) and sequencing reads from chromosome M were removed. The Peak calling of each ATAC-seq library was performed with MASC2^4^ with parameters -f BED, -g hs, -q 0.01, --nomodel, --shift 0. Peaks were merged into matrix with bedtools merge^5^. Raw reads within peaks were normalized using EdgeR’s cpm^6^. Log transformation were applied on these normalized peaks to calculate the pearson correlation among duplicates. Unique ATAC peaks for hGSC ATAC clusters were selected using DESeq2^7^, with cutoff : p value < 0.01, FDR < 0.01, log2 fold change > 1, peak average intensity > 30, and coefficient of variance < 0.2. Mouse differential ATAC-seq peaks using ATAC new cluster: cluster AB and cluster C, were identified following the principle that two clusters were compared with each other with parameter log2(fold change) > 1, false discovery rate <0.05. The ATAC-seq saturation analysis from human GSC was performed by randomly sampling samples and successively calculating the number of peaks identified within the number of samples. The self-starting non-linear mode has been used to predicate the saturation point. For ATAC-seq peak visualization, Washu Epigenome Browser was used to visualize these presentative peak regions from mouse NSC, mGSC and hGSC.

Non-linear model:

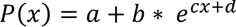

### Where P(x) represents predicted numbers of peaks, x corresponds the actual number of peaks, a,b,c and d represent the parameters for self-starting simulation

To summarize the chromatin accessibility signal per gene, the accessibility of the regions (no further than the window -1000 to +100bp from a transcriptional start site) were defined as promoter regions and the elements (located more than 3 Kbp from a TSS and no further than 500 Kbp) were represented as distal regulatory elements (DRE). The genomic annotation of ATAC-seq was performed with seven genomic features: 3’ UTR, 5’ UTR, exon, intergenic region, intron, TSS and TTS using the ChIPseeker^8^.

### NMF cluster analysis of human and mouse GSC and mouse NSC ATAC-seq

The Non-negative matrix factorization (NMF) method^9^ was used to cluster Human GSC ATAC-seq and mouse NSC and GSC ATAC-seq with nsNMF^10^. In brief, ATAC-seq peaks were ranked according to their variance from high to low. The cophenetic correlation score was calculated with cluster number 2 to 8, and used to determine the cluster numbers following the standard method^9^. Top 68000 ATAC-seq peaks (40%) from mGSC, and top 70000 (20%) ATAC-seq peaks from hGSC were used to build NMF clusters.

### Genomic segmentation analysis for the Human GSC ATAC-seq

The chromatin-state discovery and genome annotation for the ATAC-seq peaks from the human GSC ATAC-seq peaks was performed with ChromHMM^11^ by downloading the data from following dataset: GSE119755 (H3K27ac ChIP-seq); GSE121601 (H3K27ac and CTCF ChIP-seq); GSE92458 (H3K4me1 and H3K27ac ChIP-seq); GSE74557 (H3K27me3 and H3K4me3 ChIP-seq). In total, 7 chromatin status referred to Epigenomic Roadmap Consortium were defined: active promoter (H3K27ac and H3K4me3 together), active region (H3K27ac alone), inactive regions (H3K27me3 alone), insulator (CTCF), strong enhancer (H3K4me1 and H3K27ac together), weak enhancer (H3K4me1 alone) and no signal, were used to characterize the human GSC ATAC-seq peaks.

### Transcription factor motif enrichment for cluster-specific ATAC-seq peaks

Specific motifs enrichments in mGSC or hGSC cluster calculated by Deseq2^7^ was explored here. Homer^12^ vertebrate database and the coordinates of our cluster-specific peak sets were used to calculate motif occurrences^13^. TF accessibility deviation value for each sample could be obtained. The variability of the TF motifs across the whole sample set was determined by calculating the standard deviation of these. TF deviation with the threshold variability close to 1 was not significantly variable. TFs whose deviation score was positively correlated to one cluster was selected to represent each cluster based on the average of deviation score of each cluster. In order to identify the top 50 variable TFs, TFs were ranked based on their variabilities. After removing non-variable TFs, the z-score of deviation of each TF was visualized in a heatmap. Considering raw motif enrichment is not enough to predict the activity of TF, TF deviation and gene expression were combined for identifying these active TFs with the threshold p-value < 0.05 as potential mediators of the observed motif enrichment.

### ATAC-seq peak to gene linkage predictions

Putative linkage between ATAC-seq peaks and gene expression were predicted with a correlation-based approach. First, ATAC-seq peaks were annotated to the nearest genes within +/-0.5 Mbp but +/- 3 Kbp of TSS. For each pair, the Pearson correlation between the ATAC-seq peak accessibility and the gene expression level was calculated. Next, the mean and standard deviation for these correlations were calculated to represent nonspecific correlation. Then, multiple correction was performed using Benjamini-Hochberg procedure to adjust these p-values. At last, only pairs with False Discovery Rate (FDR) < 0.05 were kept.

### TFs Footprint analysis for cluster-specific ATAC-seq peaks

In previous descriptions, Tn5 transposase inserted two adaptors separated by 9bp^14^. Sequencing reads aligned files in sam format by offsetting +4/-5bp for all the reads depending on the strand of reads. A shifted base sam file converted to bam format and was sorted by samtools^15^. ATAC-seq reads for each cluster of samples (mGSC ATAC: cluster AB and cluster C; hGSC ATAC50: C1, C2 and C3) were concatenated and 200 million reads were randomly selected from cluster and merged into bam files. Then TFs footprint analysis was performed on cluster-specific regions using HINT-ATAC software. Input motif was obtained from Homer^12^ database on vertebrate. 414 motifs were tested and filtered with p-value < 0.05. The normalized read counts were centered by the motif sites around 200bp genomic region for visualizing motif footprints.

### TFs motif enrichment in cross-species analysis

After identifying the unique peaks for cluster AB and cluster C and then doing the genomic annotation for mouse peaks with ChIPseeker^8^, mouse genes were converted to their corresponding human gene. Pairs from previous peak-to-gene linking predictions were used to obtain peaks from hGSC genes. NMF method was performed on these hGSC peaks to guide the new clusters. MCO genes guided hGSC cluster was build following the same described method. For MCO promoter-guided cluster, after converting MCO gene to their relative human genes. Only promoter regions from these human genes were used to build NMF clusters. Finally, HOMER^12^ was to calculate the transcriptional factor motif enrichment from mGSC ATAC-seq cluster-specific peaks and mGSC ATAC guided hGSC peaks.

### Sphere formation assay

Established cultures from mGSC and hGSC cultures were made into single-cell suspensions by dissociating them with Acutase (Invitrogen, A1110501) and TrypLE (Thermofisher, 12563011) respectively. For primary sphere formation, 1000 cells/well were seeded in 8 replicates in a 24 well low attachment plates. After 7 days, the number of primary spheres formed for each culture were counted. For hGSCs, the primary spheres were dissociated and seeded 1000 cells/well in 8 replicates again for secondary sphere formation and counted after 7 days. Similar procedure was followed to seed and count the tertiary sphere formation of hGSC cultures.

### Proliferation analysis

Cells from hGSC cultures (5 x 10^3^ per well) were seeded (day 0) on laminin-coated coverslips in a 24 well plate using serum-free medium. The next day (day 1) 1μg/μl of BrdU (Sigma, B5002) was added to each well for 16 hours before they were fixed with 4% formaldehyde (Histolab, 02176). After fixation cells were washed with PBS and permeabilized in 2M HCl for 20 minutes, followed by the washing again with PBS. Cells on the coverslips were permeabilised in 0.2% triton X-100 with 3% Bovine serum albumin (Sigma, A7906) for 5 minutes and washed thrice with PBS. Cells were blocked in 0.2% triton X-100 solution containing 1% BSA and 5% normal goat serum (DAKO, X0907) for 1 hour.

Primary antibody again BrdU (Abcam, Ab6326) was applied overnight in a humidified chamber at 4°C. Cells were washed three times with 0.2% Triton X-100 solution for about 5 minutes each time. Secondary antibody incubation was performed for 30-60 minutes in room temperature with anti-rat Alexa 555 (1:400, Invitrogen, A21434). Lastly cells were washed three times with 0.2% Triton X-100 solution for about 5 minutes each time and mounted in Fluoromount (Sigma, F4680) with 0.1% DAPI in it. The stainings were visualised and quantified under LEICA DMi8 Fluorescent microscope. The experiment was repeated three times on consecutive passages for all the cultures.

### Extreme limiting dilution assay

Cells from hGSC cultures were dissociated and made into single-cell suspensions in serum-free medium. Cells were seeded in a 96-well low attachments plates (CLS3474-24EA, Sigma) with the seeding density ranging from 100 cells to 1 cell per well, with 10 replicates per condition. After 7-10 days, number of wells without spheres for each cell density were counted. Number of cells required to form one sphere per well was calculated by extrapolating the values of x-intercept for each culture and plotted using PRISM 7 software. The experiment was repeated three times for all cultures.

### Invasion assay

Cell spheres obtained from hGSC cultures by seeding 100/50 cells per well in ELDA experiment were used to measure the invasive capacity for each culture. Collagen gel matrix was prepared and spheres were transferred into the collagen gel matrix sandwich in a 24 well plate as described previously^16^. Pictures were taken after 10 minutes and 24 hours in Eclipse TS 100 Nikon microscope. Image J 1.52a software was used to measure the invasion area for each sphere. From each hGSC culture at least 10 spheres were analysed. The experiment was repeated three times on consecutive passages for all cultures.

### Drug response analysis in human glioblastoma cell cultures

A selected panel of 28 anti-cancer compounds were used to measure the drug sensitivity of hGSC cultures. Cells were seeded, 1000 cells/well in a poly-ornithine (Sigma, P3655) and laminin (Sigma) coated 384 well plate (Thermo fisher scientific). The following day cells were exposed to the compounds for 24 hours and were measured for their drug response after 72 hours of treatment. Cell viability was measured by using the non-clonogenic fluorometric microculture cytotoxicity assay (FMCA). Dose response curve was plotted by calculating the area under the curve values for each compound in each individual culture. The log10 fold change of the average of AUC value between cultures from paired comparison of ATAC-seq clusters were calculated using wilcoxon test and scatter plot was drawn as described previously^17^.

### In vivo xenograft analysis

All animal experiments were performed in accordance with the rules and regulations of Uppsala University and approved by the local animal ethics committee (C237/12 and C182/14). Intracranial cell transplantations of human GSC cultures were performed in neonatal NOD-SCID mice as previously described^16,18-20^. In addition to the published data 60 new mice were included (Table S8). In brief cells were dissociated in TrypLE and resuspended in DMEM/F12 medium. A volume of 2 μl cell suspension with cells ranging from 10000 till 200000 was orthotopically injected using a motorized stereotaxic injector (Stoelting CO). The coordinates measured from lambda were: anterio-posterior 1.5 mm, medio-lateral 0.7 mm and dorso-ventral 1.5 mm. Mice were monitored every second day and euthanized upon symptoms of disease. Only mice that showed disease symptoms before the endpoint of the experiment were used in the analysis.

### Quantification and Statistical analysis

Statistical analysis was performed using GraphPad PRISM 7 software or R version 3.4.0. Figures containing data from multiple repetitions of experiments were presented as mean ± SEM. For sphere-formation, ELDA, proliferation and invasion experiments students t-tests were performed. For mice and patient survival, Log-rank (Mantel-Cox) test was the statistical method used to calculate the significance in between the groups/clusters. In all the experiments the statistical significance between the groups were determined with the following p-values as *p<0.05, **p<0.01, ***p<0.001, and ****p<0.0001.

